# *KEAP1* mutations activate the NRF2 pathway to drive cell growth and migration, and attenuate drug response in thyroid cancer

**DOI:** 10.1101/2025.08.13.670155

**Authors:** Nicholas E. Bambach, Julio C. Ricarte-Filho, Erin R. Reichenberger, Christian Hernandez-Padilla, Kyle Hinkle, Amber Isaza, Andrew J. Bauer, Aime T. Franco

## Abstract

The KEAP1/NRF2 pathway, a major regulator of the cellular oxidative stress response, is frequently activated in human cancers. Often mediated by loss-of-function mutations in *KEAP1*, this activation causes increased NRF2 transcriptional activity and constitutive activation of the antioxidant response. While *KEAP1* mutations have been well documented in various cancers, their presence and role in thyroid carcinoma have remained largely unexplored. In this study, we sequenced pediatric thyroid tumors and analyzed publicly available datasets, identifying 81 *KEAP1* mutations in tumors across a range of histologies. In these tumors, we further identified frequent biallelic loss of *KEAP1* via 19p13.2 loss of heterozygosity (LOH). MAPK-activating alterations were found in a subset of *KEAP1*-mutant cases, but they were mutually exclusive with 19p13.2 LOH. Transcriptome analysis also revealed significant activation of the NRF2 pathway in *KEAP1-*mutant tumors. Four additional cases with similar transcriptional profiles but lacking mutational data were identified, likely representing putative *KEAP1* mutants. Using *in vitro* cell line models, we then profiled the functional consequences of *KEAP1* knockout in cells with and without known driver alterations. In these models, we show that *KEAP1* loss leads to an NRF2-dependent upregulation of *AKR1C3, GCLC, NQO1*, along with increased proliferation and migration, irrespective of MAPK mutational status. We also demonstrate that loss of *KEAP1* reduced sensitivity of *RET* fusion-positive cells to selpercatinib, consistent with previous reports that these alterations promote drug resistance in other malignancies. In this report, we comprehensively profile *KEAP1* mutations in thyroid tumors, showing they are more prevalent and functionally significant than previously recognized. These findings position *KEAP1* mutations as potential novel oncogenic drivers in thyroid cancer and support the integration of KEAP1/NRF2 pathway profiling into future studies and clinical frameworks.

## INTRODUCTION

The kelch like ECH associated protein 1 (KEAP1)/nuclear factor erythroid 2-related factor 2 (NRF2) pathway is the master regulator of cellular defense against chemical toxicity and oxidative stress. Under normal physiological conditions, reactive oxygen species (ROS) are produced and play critical regulatory roles in immune function, inflammation, cell division, and stress response (1). However, uncontrolled production of these oxidant species impairs metabolic function, causes toxicity, and contributes to the pathophysiology of disease, including tumorigenesis (2, 3). Accumulation of ROS is therefore counterbalanced by a highly regulated and complex antioxidant response, primarily mediated by the KEAP1-NRF2 pathway.

KEAP1, encoded on chromosome 19p13.2, is a Broad complex Tramtrack and Bric à Brac (BTB)-Kelch protein that functions as a negative regulator of NRF2 by facilitating its ubiquitination by the CUL3-based E3 ubiquitin ligase complex (4, 5). Under normal conditions, KEAP1 sequesters NRF2 in the cytoplasm, allowing for its polyubiquitination and subsequent degradation by the 26S proteasome (6). This maintains low basal levels of NRF2 and avoids inappropriate activation of the chemoprotective response (7). Under oxidative and electrophilic stress, however, radicals and electrophiles covalently modify reactive cysteine residues on KEAP1, diminishing the KEAP1-CUL3 complex’s ubiquitin ligase activity and stabilizing NRF2 (8–13). As it accumulates, NRF2 translocates to the nucleus, binds the antioxidant response element (ARE), a *cis-*acting enhancer sequence in the 5’-promoter region of cytoprotective genes, and induces transcription (10, 14–16).

While activation of NRF2 and its target genes has been shown to prevent cancer by eliminating carcinogenic oxidants and toxins, its constitutive activation can also drive tumorigenesis (17–20). In fact, NRF2 overexpression has been found to promote malignancy, tumorigenesis, proliferation, and migration, as well as confer resistance to cytotoxic agents (21–28). Cancers frequently achieve overexpression of NRF2 and these resulting phenotypes via somatic loss-of-function mutations in *KEAP1*. This has been well documented in renal, ovarian, esophageal, head and neck, gallbladder, and prostate cancer, and occurs at particularly high frequency in non-small cell lung cancer (18, 19, 29–33). Aberrant activation of NRF2 via *KEAP1* mutation is also associated with poor clinicopathological outcomes, including low-response rate to platinum-based chemotherapy, increased metastasis (long distance, lymph node, and tumor node), poor overall survival rate, and poor progression-free survival (34, 35). In addition to somatic mutations, epigenetic silencing of *KEAP1* through promoter hypermethylation has also been reported in colorectal cancer, breast cancer, lung cancer, and kidney cancer (36–41). This hypermethylation is associated with increased NRF2 transcriptional activity, increased disease progression, higher mortality rate, and decreased response to chemotherapeutics.

In thyroid cancer, mutations that activate the MAPK pathway are the most common oncogenic drivers. However, approximately 10% of thyroid cancers have unidentified driver mutations (42, 43). Due to the prevalence of *KEAP1* alterations in other malignancies, we sought to determine whether these mutations could represent an alternative driver of thyroid cancer, accounting for these unidentified mutations, or alternatively cooperate with other MAPK-activating mutations to further drive pathogenesis of disease. Limited evidence has implicated the KEAP1/NRF2 pathway in thyroid tumorigenesis; however, this connection remains incompletely characterized. In rare cases of familial non-toxic multinodular goiter, *KEAP1* mutations have been identified and linked to increased NRF2 accumulation and transcriptional activity (44, 45). In papillary thyroid carcinoma (PTC), a handful of *KEAP1* alterations have been identified and associated with increased nuclear NRF2 expression, correlating the mutations with NRF2 stabilization (46). A retrospective immunohistochemical analysis also identified high NRF2 levels in PTC relative to benign lesions, indicating NRF2 pathway activation as a potential contributor to thyroid tumorigenesis (47). Additionally, an analysis of PTC samples from The Cancer Genome Atlas (TCGA) revealed epigenetic silencing of *KEAP1* through promoter hypermethylation, further linking KEAP1/NRF2 pathway dysregulation to thyroid carcinogenesis (48).

While these previous reports identify *KEAP1* mutations and implicate the KEAP1-NRF2 pathway in thyroid cancer, this connection has remained incompletely characterized. To our knowledge, this is the first study to comprehensively report the presence of inactivating *KEAP1* mutations across all thyroid cancer subtypes. We characterize these mutations in the broader genomic landscape and identify for the first time a frequent genetic mechanism for biallelic *KEAP1* loss. This study is also the first to comprehensively explore the functional consequences *KEAP1* loss in thyroid carcinoma. We investigate the impact of *KEAP1* loss on NRF2 pathway activation, as well as its effects on cell growth, migration, and drug response. Our findings indicate that *KEAP1* mutations may represent a novel driver of thyroid tumorigenesis, as observed in other malignancies.

## MATERIALS AND METHODS

### Patient samples and targeted DNA/RNA sequencing

Patient samples were collected at the time of surgery and stored under the infrastructure of the Child and Adolescent Thyroid Consortium (CATC) Biorepository (IRB# 17-014224 and IRB# 20-018240). Histopathologic results, tumor staging, and demographic information including age and sex were collected from each patient and deidentified within the CATC Central RedCap data repository (49).

This study involved human-derived biospecimens and was approved by the CHOP Institutional Review Board (IRB) with two arms: (a) for patients who had surgery before the time of initial IRB approval, a waiver of consent was approved as these were existing biospecimens from previous surgeries; and (b) for patients who had surgery after the time of initial IRB approval, enrollment is considered prospective and thus consent and assent were obtained.

Tumors were sequenced using the CHOP Division of Genomic Diagnostics’ Comprehensive Solid Tumor Panel (CSTP), which evaluates sequence and copy number (CN) alterations in 238 cancer associated genes, including *KEAP1*. mRNA expression was profiled using the HTG EdgeSeq Oncology Biomarker Panel (OBP), as previously described (50).

### Data sources

Genetic data and clinical information for additional cases were collected from the following studies: AACR Project GENIE, Global Anaplastic Thyroid Cancer Initiative (GATCI), ICGC-THCA, MSK-IMPACT, MSK-MET, TCGA-THCA, Danilovic et al., Eszlinger et al., Ganly et al., Gopal et al., Kunstman et al., Landa et al., Lee et al., Nishihara et al., Pozdeyev et al., and Teshiba et al. (44–46, 51–62). Data from these studies were accessed and downloaded from publicly available databases including the Genomic Data Commons (GDC), cBioPortal, and COSMIC, or from the original publication.

For applicable studies, segmented CN data was downloaded and visualized in the Integrative Genomics Viewer (IGV). A log_2_(CN) ≤ −0.2 threshold was set to designate cases with heterozygous segment loss at the relevant locus. mRNA expression data was downloaded and extracted from the Gene Expression Omnibus (GEO) using the GEOquery package (v2.74.0) in RStudio (v4.4.2).

### Generation of clonal *KEAP1* knock-out cell lines

TPC1 and SDAR2 cells were transfected with three distinct sgRNAs (Synthego) targeting sites in exon 2 of *KEAP1* using Lipofectamine 3000 (Thermo Fisher Scientific) according to the manufacturer’s instructions. Ten experimental and two control transfections were performed per cell line. Sanger sequencing and Inference of CRISPR Edits (ICE, Synthego) were used to evaluate knock-out efficiency. Cells were then sorted using a FACSMelody Cell Sorter (BD Biosciences) and seeded at a density of 1 cell per well in a 96 well plate. Cells were cultured in growth medium supplemented with 20% FBS, 1% penicillin-streptomycin, 1X piperacillin-ciprofloxacin, and 1X amphotericin B, screened for mRNA expression of NRF2 target genes, and Sanger sequenced to confirm successful knockout.

### DNA extraction and Sanger sequencing

DNA was isolated using the Quick-DNA Miniprep Kit (Zymo Research, Irvine, CA) according to the manufacturer’s instructions. 50ng of DNA was PCR amplified, and the resulting product was purified with the ZR DNA Sequencing Clean-up Kit (Zymo Research). 10ng of the purified PCR product was Sanger sequenced using the forward and reverse primers. The sequence was extracted from the chromatogram using 4Peaks (Nucleobytes) and sequence alignment was performed using the Basic Local Alignment Search Tool (BLAST, NCBI). Primer information is listed in Supplementary Materials and Methods.

### RNA isolation and RT-qPCR

RNA was extracted using TRIzol Reagent (Thermo Fisher Scientific) and the Direct-zol RNA Miniprep Kit (Zymo Research). 1000ng of RNA was reverse transcribed using the Verso cDNA Synthesis Kit and PCR reactions were performed using PowerTrack SYBR Green Master Mix (Applied Biosystems). Gene expression was measured in triplicate using the Quant Studio Real-Time PCR System (Applied Biosystems). β-actin (ACTB) was used to normalize expression of all samples. Primer information is listed in Supplementary Materials and Methods.

### Immunofluorescence imaging

Cells were fixed and permeabilized with a solution of 4% paraformaldehyde, 5% (w/v) sucrose and 0.1% Triton X-100. The fixed cells were blocked with 3% BSA for 1 hour at room temperature, incubated with the primary antibody overnight at 4°C, and incubated with the secondary antibody for 1 hour at room temperature. Nuclei were counterstained with 4’,6-diamidino-2-phenylindole (DAPI, 1µg/mL) for 15 minutes. Cells were imaged at 20X via confocal microscopy using the ZEISS LSM980. NRF2 nuclear intensity was quantitated using ImageJ. The following primary antibodies were used: NRF2 (1:200, Cell Signaling), Rhodamine Phalloidin-TRITC (1:400, Invitrogen). The following secondary antibody was used: Alexa Fluor™ Plus 555 Goat anti-Rabbit IgG (H+L) (1:500, Invitrogen).

### Western blotting

Cells were lysed with radioimmunoprecipitation assay (RIPA) buffer supplemented with 1X Halt Protease and Phosphatase Inhibitor Cocktail and phenylmethylsulfonyl fluoride (PMSF) (Thermo Scientific). Protein concentration was determined using bicinchoninic acid (BCA) assay and 10µg of protein was loaded on a 10% polyacrylamide gel and resolved via SDS-PAGE. Proteins were transferred to a PVDF membrane (Millipore) and blocked with 5% BSA for 1 hour at room temperature. Membranes were incubated with the primary antibody overnight at 4°C, secondary antibody for 1 hour at room temperature, and were visualized using SuperSignal™ West Femto Maximum Sensitivity Substrate (Thermo Scientific). The following primary antibodies were used: KEAP1 (1:1000, Cell Signaling), β-actin (1:5000, Cell Signaling). The following secondary antibody was used: HRP-linked Goat anti-Rabbit IgG (1:5000, Cell Signaling).

### Cell proliferation assays

Growth curves were performed using the CellTiter-Glo luminescent assay. TPC1 cells (1500 cells/well in 5% FBS) and SDAR2 cells (1500 cells/well in 10% FBS) were seeded in 96-well plates. At the indicated time points, CellTiter-Glo reagent was added to each well at a 1:1 ratio with culture medium and the luminescence was measured using a Varioskan LUX microplate reader (Thermo Scientific). Luminescence was averaged across 10 replicates.

### Transwell migration assays

Cells (10^5^/well) were seeded in the upper chamber of a 24-well transwell chamber (8µm pore size, Corning) in the recommended FBS-free medium containing 0.1% BSA. Complete medium containing 10% FBS was added to the bottom chamber of the insert as a chemoattractant. After 16 hours of incubation, non-migrating cells were removed from the upper chamber using a cotton swab and the remaining cells were fixed and stained using 70% ethanol and 0.5% (w/v) crystal violet (Fisher Chemical). The number of migrated cells was counted from five randomly selected fields.

### Dose-response curves

TPC1 cells (1500/well in 5% FBS) were seeded in 96-well plates. After 24 hours, cells were dosed with the indicated concentrations of selpercatinib diluted in culture medium. Cells were cultured for 72 hours and nuclei were stained with NucBlue Live ReadyProbes Reagent (Invitrogen). The stained nuclei were then imaged with the EVOS M7000 and quantified using ImageJ. Cell counts were normalized to percent of control and the IC_50_ was calculated using GraphPad’s four parameter logistic regression.

### Colony formation assays

TPC1 cells (500/well in 5% FBS) were seeded at single cell density in 6 well plates. After 24 hours, cells were dosed with the indicated concentration of drug diluted in culture medium. Drug treatments were refreshed every 3 days. After 12 days in culture, colonies were fixed and stained using 100% ice-cold methanol and 0.5% (w/v) crystal violet (Fisher Chemical). Wells were imaged then crystal violet was eluted using 10% acetic acid. Absorbance was recorded at 595 nm on a Varioskan LUX microplate reader (Thermo Scientific).

### Statistical analyses

All statistical tests were performed with GraphPad Prism 10 (v10.4.1) or in RStudio (v4.4.2). Fisher’s Exact Test, one-way ANOVA, and two-way ANOVA were all conducted and a two-sided p-value < 0.05 was considered statistically significant.

## RESULTS

### *KEAP1* mutations occur in both pediatric and adult thyroid tumors

Mutations in *KEAP1* have previously been reported in various pediatric and adult cancers but have not been widely reported or well characterized in the context of thyroid tumors. To assess the prevalence of these mutations in pediatric tumors, we sequenced samples from the CHOP Biorepository (n = 191) using the CSTP and identified five papillary carcinomas and one benign multinodular goiter harboring *KEAP1* mutations (Figure 1, Table S1). These six cases represented 3.14% of the total tumors in our biorepository and included four females with a mean age at surgery of 15.19 (SD = 2.23). Four of the *KEAP1* mutations were missense, while two were truncating. We next sought to determine if *KEAP1* mutations are also present in non-pediatric cases. We examined sequencing data from publicly available datasets and published literature and identified an additional 75 tumor samples and two thyroid cancer cell lines with *KEAP1* mutations (Figure 1, Table S1). The identified cell lines were previously established from primary tumors and not reported in tumor databases, thus representing independent cases. These cases included 36 females with an average age at surgery of 58.04 (SD = 13.63). Of these cases, 51 had missense mutations, 23 had truncating mutations, and one had an in-frame deletion.

**Figure 1:**
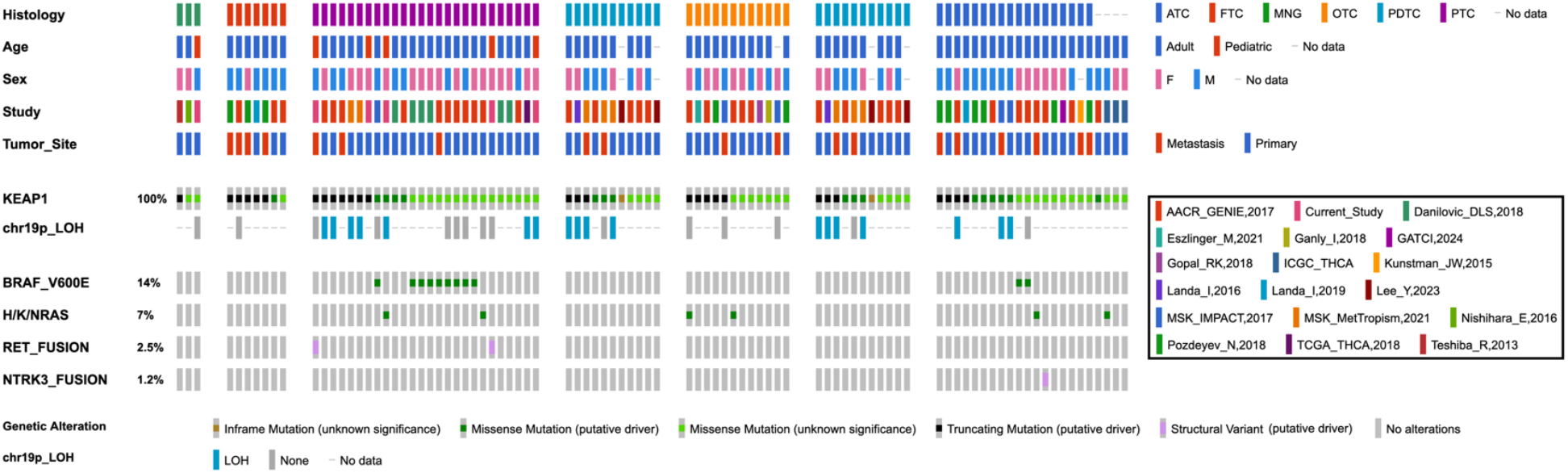
*KEAP1* mutations occur in both pediatric and adult thyroid tumors. Oncoprint detailing 81 *KEAP1*-mutant thyroid tumors, including cases from the current study and publicly available datasets. Clinicopathologic features include histology, age, sex, tumor origin, type of *KEAP1* variant, chromosome 19p13.2 loss, and MAPK alteration. The predicted consequence of each *KEAP1* mutation is annotated based on its OncoKB prediction. Abbreviations: ATC = anaplastic thyroid carcinoma, FTC = follicular thyroid carcinoma, MNG = multinodular goiter, OTC = oncocytic thyroid carcinoma, PDTC = poorly-differentiated thyroid carcinoma, PTC = papillary thyroid carcinoma, LOH = loss of heterozygosity.

Between our CHOP cohort and publicly mined data, we identified 81 total cases representing a range of histologies: 3 multi-nodular goiters (MNGs) (3.70%), 7 follicular thyroid carcinomas (FTCs) (8.64%), 26 papillary thyroid carcinomas (PTCs) (32.1%), 12 oncocytic thyroid carcinomas (OTCs) (14.8%), 11 poorly-differentiated thyroid carcinomas (PDTCs) (13.6%), 18 anaplastic thyroid carcinomas (ATCs) (22.2%), and 4 cases with unspecified histologies (4.94%). Mutation mapping of these 81 *KEAP1* mutations revealed that variants were distributed across all annotated domains of *KEAP1*, showing no evidence of clustering within a specific region (Figure S1). Additionally, in cases where mutational status could be determined, 58 (92.1%) mutations were confirmed somatic, while 5 (7.94%) were confirmed germline. These five germline cases included three from our CHOP cohort which were confirmed to be germline variants via Sanger sequencing or whole exome sequencing of adjacent normal tissue (Figure S2). For the remaining 18 cases, mutation status was unable to be evaluated due to limited data in publicly available cases or absence of paired normal tissue in the CHOP cohort.

We next examined CN segment data to determine if any *KEAP1-*mutant cases had concurrently lost their remaining wild-type allele. In 14 (50%) of cases with reported CN data, we identified loss of heterozygosity (LOH) at chromosome 19p13.2, indicating biallelic loss of wild-type *KEAP1*. In 24.7% of the *KEAP1-*mutant tumors, we also identified several concurrent MAPK-activating mutations: BRAF p.V600E, KRAS p.G12V/D, NRAS p.Q61R/K, *RET*, or *ETV6::NTRK3* fusions. Notably, these MAPK alterations were mutually exclusive with 19p13.2 LOH in *KEAP1*-mutant tumors (OR = 0.049, *p* = 0.0044). Figure 1 displays an oncoprint plot detailing all cases harboring *KEAP1* mutations, 19p13.2 loss, concurrent MAPK alteration, and associated demographic and histopathologic features. Table S1 provides detailed molecular and demographic information for each sample.

### *KEAP1* mutations activate the NRF2 pathway in thyroid tumors

To determine whether *KEAP1* mutations correlated with NRF2 pathway activation, we profiled mRNA expression of five canonical NRF2 target genes: *AKR1C3, GCLC, GSR, NQO1*, and *TXNRD1*. Six *KEAP1-*mutant tumors—four PTCs, one OTC, and one PDTC—had associated mRNA expression data. Four of these cases had missense *KEAP1* mutations, while the others had truncating alterations (Figure 2A). In addition, we identified one FTC-derived cell line (described below) with a truncating *KEAP1* mutation. Consistent with our prior observations, two of the six tumor cases exhibited concurrent MAPK pathway alterations: one carried a *CCDC186::RET* fusion, while the other had a KRAS p.G12V point mutation (Table S1). Notably, three tumors also harbored concurrent 19p13.2 LOH. In five of the six cases, we found significant overexpression of canonical NRF2 target genes (Figures 2B-E). The only case lacking NRF2 pathway activation carried a germline KEAP1 p.C226P missense mutation, which co-occurred with the *CCDC186::RET* fusion and was computationally predicted to be tolerated via SIFT (Figure S2, Table S1).

**Figure 2:**
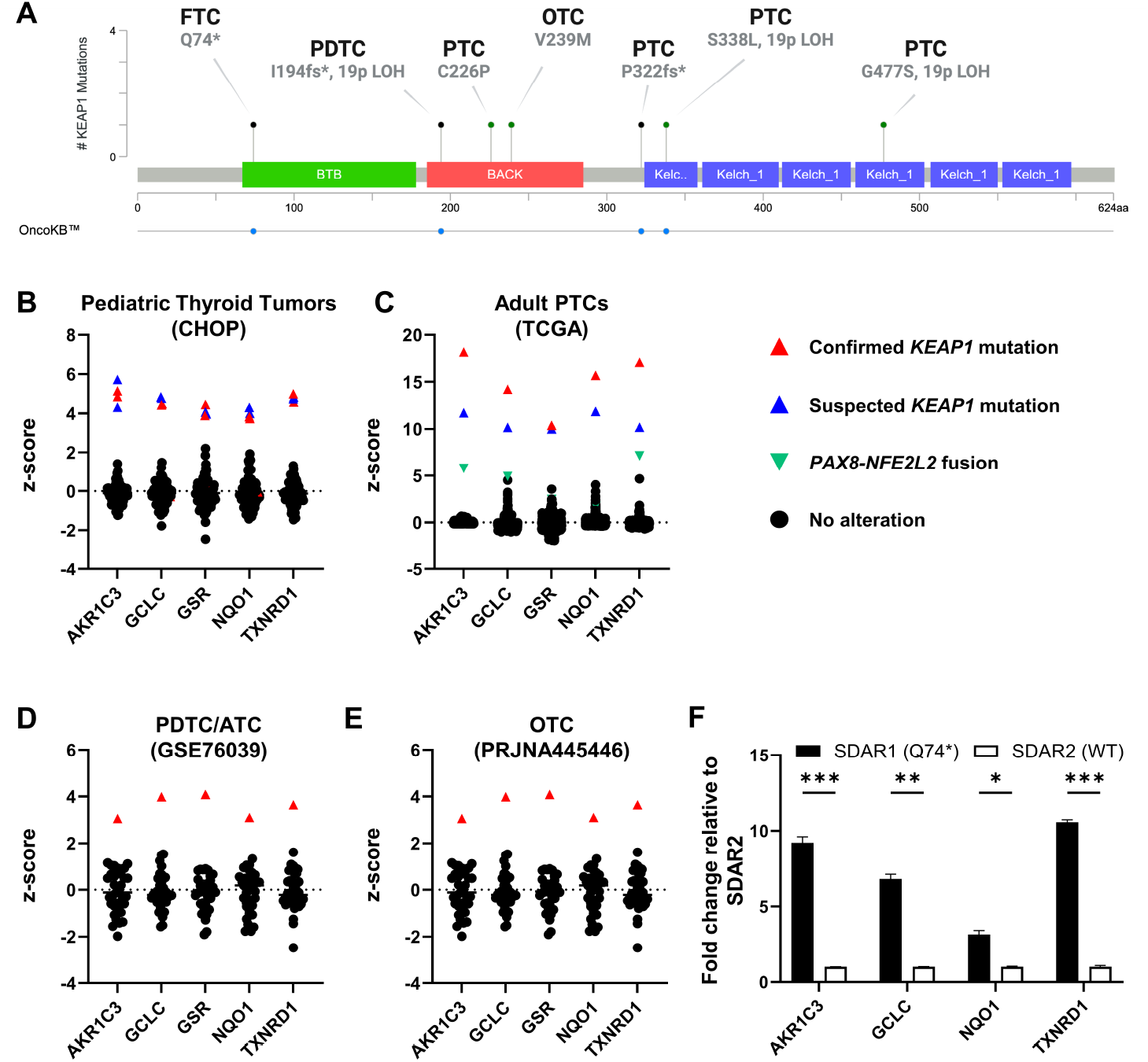
*KEAP1* mutations activate the NRF2 pathway in both pediatric and adult thyroid tumors. **(A)** Lollipop plot detailing histology and *KEAP1* alterations in tumor samples and FTC cell line with paired RNA profiling and mutational analysis. **(B-E)** RNA expression of NRF2 target genes: *AKR1C3, GCLC, GSR, NQO1*, and *TXNRD1* in **(B)** pediatric PTC/adenoma CHOP samples (n = 98), **(C)** adult PTC TCGA-THCA samples (n = 481), **(D)** adult PDTC/ATC samples from Landa, et al., 2016 (GSE76039) (n = 37), and **(E)** adult OTC samples from Ganly et al., 2018 (PRJNA445446) (n = 49). Samples of interest are colored based on their known or suspected KEAP1/NRF2 pathway alterations. Z-scores were calculated relative to all samples in the cohort. **(F)** RT-qPCR expression of NRF2 target genes *AKR1C3, GCLC, NQO1*, and *TXNRD1* in SDAR1 (KEAP1^mut^) and SDAR2 (KEAP1^wt^) cell lines. * p < 0.05, ** p < 0.01, *** p < 0.001. Statistics were performed using a one-way ANOVA.

We also identified one PTC and one follicular adenoma in our CHOP cohort for which we had RNA sequencing data which demonstrated increased expression of the same NRF2 target genes (Figure 2B). There was not remaining sample available to complete *KEAP1* mutational analysis, but the expression profiles support a loss-of-function *KEAP1* mutation or other NRF2-activating alteration. We further observed two PTCs in the TCGA cohort that also had significant overexpression of *AKR1C3, GCLC, GSR, NQO1*, and *TXNRD1* (Figure 2C). One case harbored a *PAX8::NFE2L2* fusion, causing markedly increased expression of *NFE2L2*, the gene encoding NRF2, resulting in a similar functional effect as *KEAP1* loss (Figure S3). The second case lacked mutational data but exhibited CN loss of chromosome 19p13.2. This, in combination with its expression profile, supports a putative loss-of-function *KEAP1* mutation.

In addition to patient samples, we analyzed two independent FTC cell lines, SDAR1 and SDAR2. These cell lines originated from the same patient, but from different sites: a primary tumor and a neck metastasis, respectively. It has been previously reported that SDAR1 harbors a homozygous p.Q74* mutation in *KEAP1*, while SDAR2 does not (43). We sequenced these cell lines to evaluate their mutational status and confirmed that SDAR1 indeed carries a homozygous p.Q74* mutation, while SDAR2 is wild type for *KEAP1* (Figure S4). Consistent with *KEAP1* loss and NRF2 pathway activation, SDAR1 showed increased expression of *AKR1C3, GCLC, NQO1*, and *TXNRD1* compared to SDAR2 (Figure 2F).

### Knockout of *KEAP1* causes NRF2 pathway activation in PTC and FTC cells

Since we determined that *KEAP1* mutations occur in tumors irrespective of MAPK driver status, we next sought to determine the effects of *KEAP1* loss in cell lines with and without MAPK pathway alterations. Using the PTC-derived cell line TPC1, which carries a *CCDC6::RET* fusion, we knocked out *KEAP1* to generate two clonal cell lines. Successful knockout was confirmed via Sanger sequencing and Western blotting (Figure S5). Since NRF2 transcriptional activity is dependent on its nuclear localization, we performed immunofluorescence to determine the localization of NRF2. We showed that both knockout cell lines exhibited increased levels of nuclear-localized NRF2 protein relative to the control cell line (Figures 3A, B). To further confirm increased NRF2 transcriptional activity, we evaluated the expression of the canonical NRF2 target genes *AKR1C3, GCLC, NQO1*, and *TXNRD1*, and found significant upregulation in response to *KEAP1* knockout (Figure 3C). To ensure that induction of these genes was NRF2-dependent, we treated cells with the NRF2 inhibitor ML385 and observed that expression levels of these target genes decreased in the knockout cells, with little effect on the control cell line (Figure 3C).

**Figure 3:**
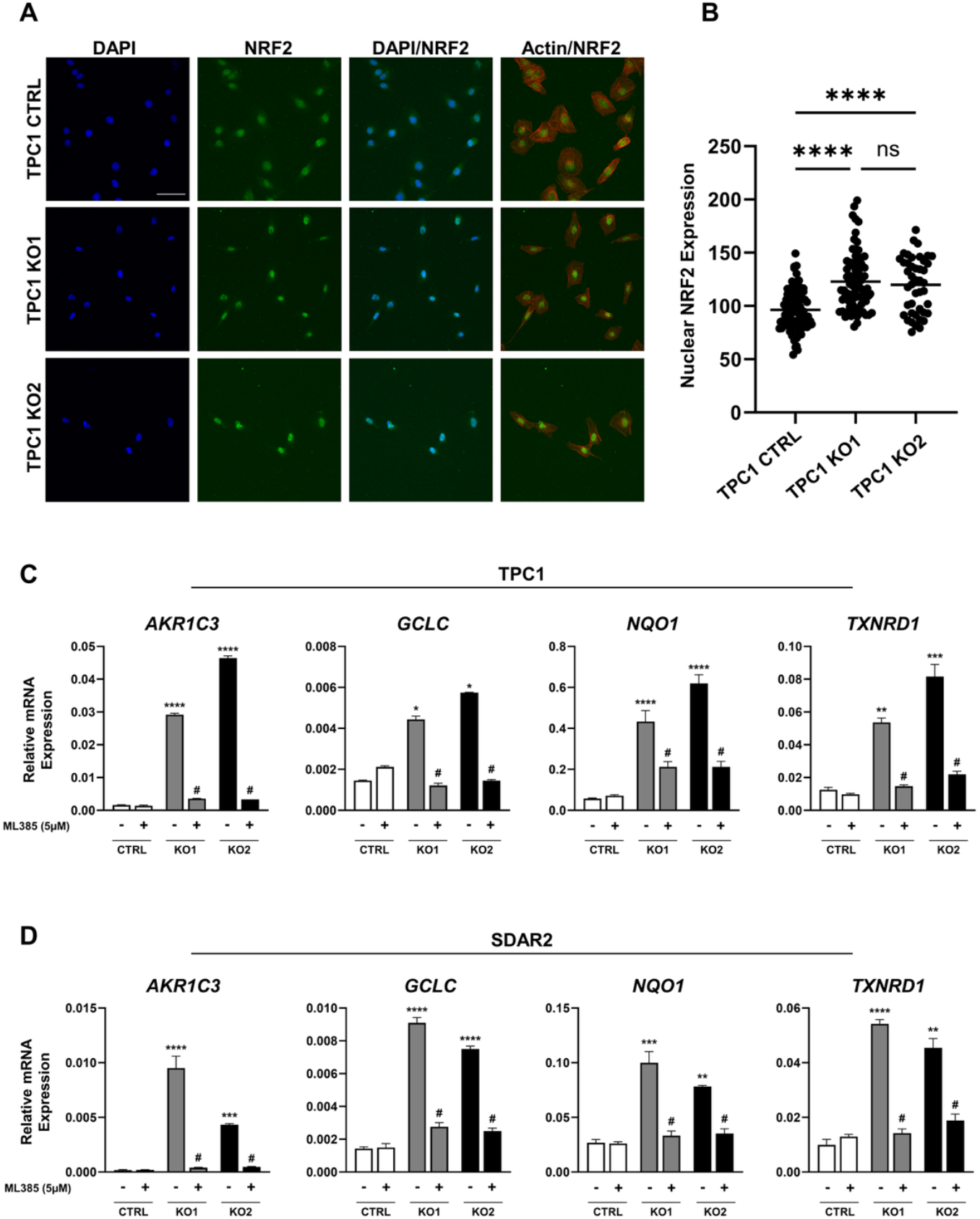
Knockout of *KEAP1* causes NRF2 pathway activation in PTC and FTC cells. **(A)** Representative immunofluorescence images of NRF2 (green), F-actin (red), and nuclei (DAPI, blue) in TPC1 control and *KEAP1* knockout clones (KO1 and KO2), showing increased NRF2 nuclear localization upon *KEAP1* knockout (scale bar = 20µm). **(B)** Quantitation of NRF2 nuclear intensity in TPC1 control (n = 84), TPC1 KO1 (n = 79), and TPC1 KO2 (n = 44) cells. **(C, D)** RT-qPCR expression of NRF2 target genes: *AKR1C3, GCLC, NQO1*, and *TXNRD1* in **(C)** TPC1 *KEAP1* knockout clones in the presence and absence of 5µM ML385 and **(D)** SDAR2 *KEAP1* knockout clones in the presence and absence of 5µM ML385, showing NRF2 dependent upregulation of target genes in knockout cell lines (black/gray bars) relative to control cell lines (white bars). All experiments were repeated in triplicate. * p < 0.05, ** p < 0.01, *** p < 0.001, **** p < 0.0001 versus the control cell line in the absence of ML385. # p < 0.001 versus the respective knockout cell line in the absence of ML385. Statistics were performed using a two-way ANOVA test.

We next examined the effects of *KEAP1* loss in cells lacking a MAPK driver. Using SDAR2, an FTC cell line with no known driver mutation, we knocked out *KEAP1* and generated two clonal cell lines. We again confirmed successful knockout via Sanger sequencing and Western blotting (Figure S5). SDAR2 cell lines did not tolerate the serum starvation necessary for the assessment of NRF2 protein localization via immunofluorescence. However, we were still able to determine NRF2 transcriptional activity by evaluating expression of *AKR1C3, GCLC, NQO1*, and *TXNRD1*. We found significant upregulation of these genes in the knockout clones relative to the control cell line (Figure 3D). To confirm this expression pattern was NRF2-dependent, we again treated cells with ML385 and saw a marked reduction in expression of the target genes in the knockout cell lines, while control cells remained largely unaffected (Figure 3D).

### Loss of *KEAP1* promotes proliferation and migration in PTC and FTC cells

We next sought to assess the functional consequences of *KEAP1* loss in cell lines with and without MAPK driver alterations by evaluating proliferation and migration. Using TPC1 *KEAP1* knockout clones, we first evaluated proliferation utilizing the CellTiter-Glo Assay. We found that loss of *KEAP1* increased growth, even in the presence of a MAPK driver mutation (Figure 4A). To then determine if this effect persisted in cells without a known driver alteration, we performed growth assays using SDAR2 *KEAP1* knockout clones and observed a similar increase in proliferation relative to control cells (Figure 4B). We next assessed whether *KEAP1* loss affects migration in MAPK-mutant and MAPK-wildtype cells by performing transwell migration assays. We observed that both TPC1 and SDAR2 cells exhibited increased cell migration upon *KEAP1* knockout, suggesting that *KEAP1* loss promotes cell migration independent of MAPK mutational status (Figures 4C-4F).

**Figure 4:**
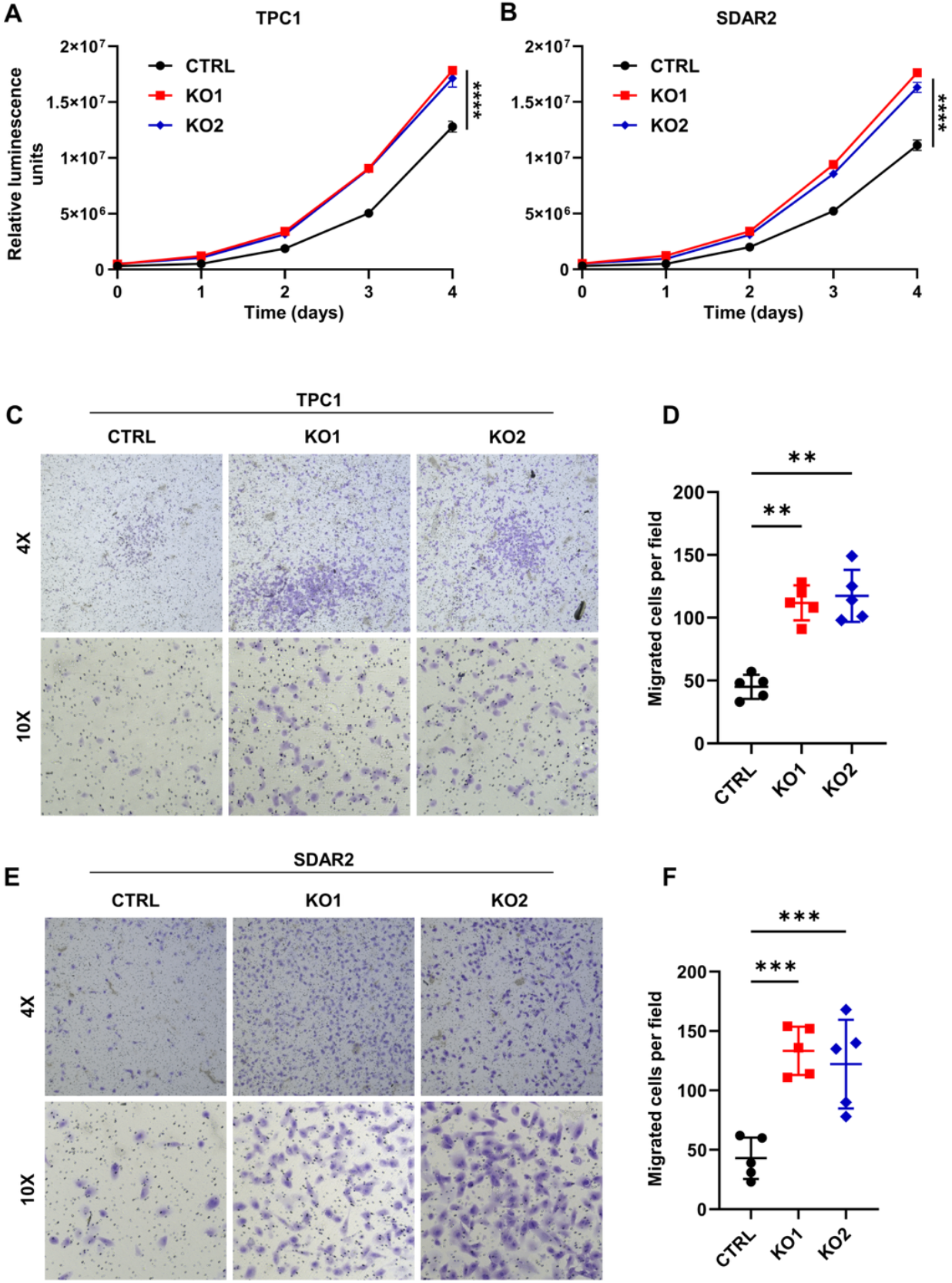
Loss of *KEAP1* stimulates proliferation and migration in PTC and FTC cells. Growth curves of **(A)** TPC1 control and *KEAP1* knockout cell lines (KO1 and KO2) and **(B)** SDAR2 control and *KEAP1* knockout cell lines (KO1 and KO2), showing increased proliferation in the knockout clones relative to controls. **(C-F)** Transwell migration assays of **(C)** TPC1 control and *KEAP1* knockout clones (KO1 and KO2) and **(E)** SDAR2 control and *KEAP1* knockout clones (KO1 and KO2) with relative quantifications **(D, F)** after crystal violet staining, showing increased migration upon *KEAP1* loss. The number of migrated cells was averaged from five randomly selected 10X fields. ** p < 0.01, *** p < 0.001, **** p < 0.0001. Statistics were performed using a one-way ANOVA test.

### *KEAP1* loss attenuates the response of *RET* fusion-positive cells to targeted receptor tyrosine kinase inhibition

The KEAP1/NRF2 pathway regulates the transcriptional output of antioxidant and drug-metabolizing genes and has been implicated in drug resistance across multiple cancers (10, 28). We therefore investigated whether *KEAP1* loss modulates the sensitivity of thyroid cancer cells to targeted *RET* inhibition. Selpercatinib is a selective *RET* tyrosine kinase inhibitor approved for use in *RET* fusion-positive thyroid cancers (63). Previous studies have shown that TPC1, which harbors a *CCDC6::RET* fusion, is sensitive to selpercatinib (64–66). We tested the efficacy of selpercatinib in TPC1 cells with and without knockout of *KEAP1*. We found that selpercatinib inhibited TPC1 parental cells with an IC_50_ of 6.2 nM, consistent with prior reports (64–66). Loss of *KEAP1* significantly reduced sensitivity to selpercatinib, with IC_50_ values of 22.4 and 25.7 nM (Figure 5A). In the *KEAP1* knockout cells, we observed a plateau in response to selpercatinib from 50-1000nM, indicating reduced sensitivity even at high inhibitor concentrations. To determine whether this effect persisted during extended selpercatinib exposure, we performed 12-day colony formation assays at high inhibitor concentrations. The TPC1 *KEAP1* knockout cells remained less sensitive to selpercatinib than control cells, even after extended exposure (Figure 5B and 5C).

**Figure 5:**
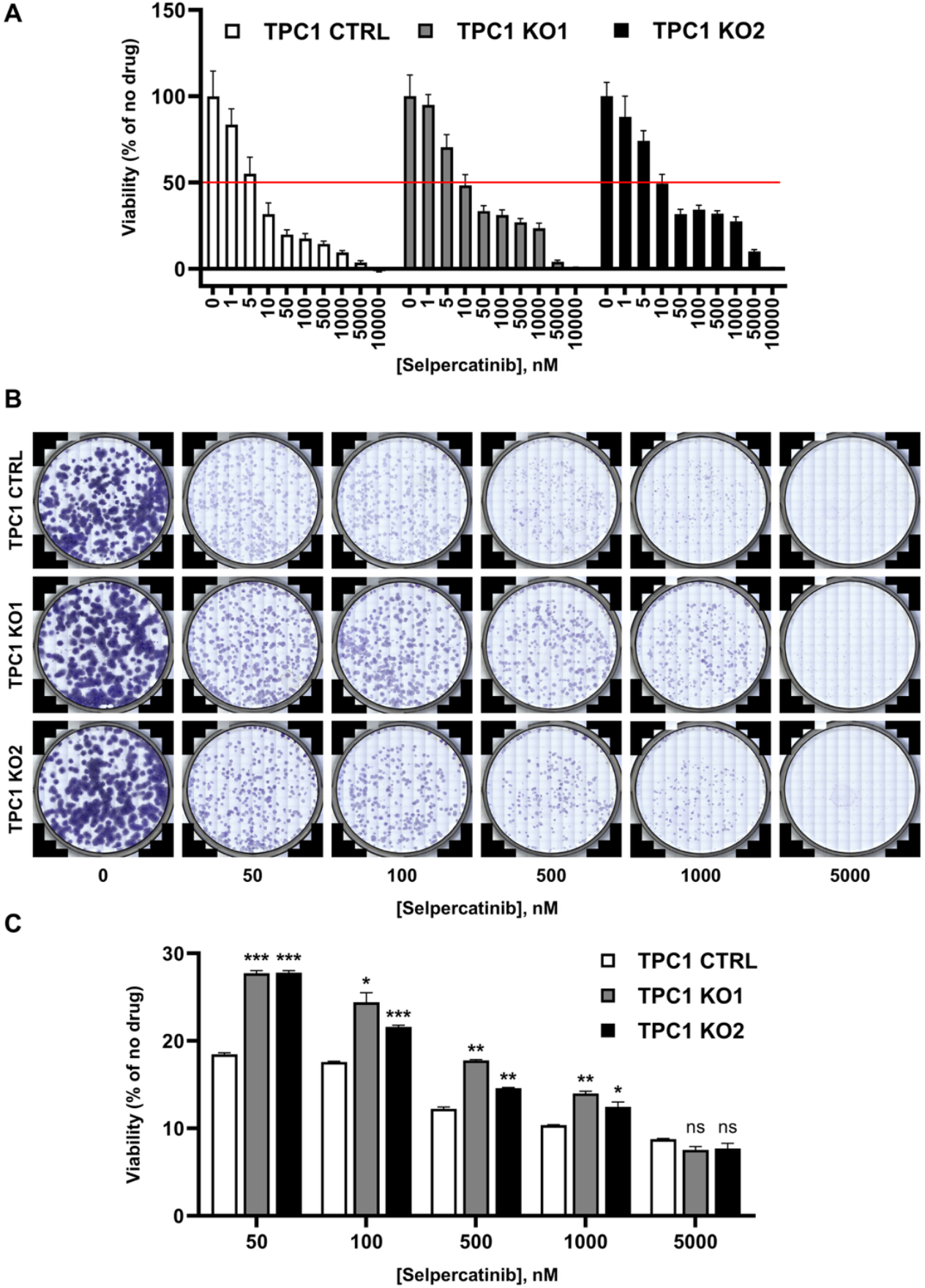
Loss of *KEAP1* desensitizes *RET* fusion-positive cells to targeted receptor tyrosine kinase inhibition. **(A)** Dose-response graph showing TPC1 control and TPC1 *KEAP1* knockout clones’ response to varying concentrations of selpercatinib after 3 days of treatment. Cell numbers were counted after nuclei staining, and percent viability was calculated relative to the 0nM condition for each cell line. **(B)** 12-day colony formation assay showing clonogenic potential of TPC1 control and TPC1 *KEAP1* knockout cell lines in the presence of high-dose selpercatinib. Colonies were stained with crystal violet and each panel is an entire well of a 6-well dish. **(C)** Quantitation via crystal violet elution. OD measurements were taken, and percent viability was calculated relative to the 0nM condition for each cell line. All experiments were performed in triplicate. * p < 0.05, ** p < 0.01, *** p < 0.001 versus control cell line at each concentration of selpercatinib. Statistics were performed using a one-way ANOVA test.

## DSICUSSION

While *KEAP1* mutations have been widely reported and studied in malignancies such as non-small cell lung cancer, their presence and role in thyroid cancer have been largely overlooked (67, 68). Although a few *KEAP1* mutations have been reported in thyroid tumors, these studies lacked systematic evaluation of NRF2 pathway activation and *in vitro* modeling of mutation consequence (44–46). Having observed *KEAP1* mutations in a number of pediatric thyroid cancer cases, we sought to determine if they represent a novel driver mutation of thyroid tumorigenesis, as has been reported in other malignancies. Here, we report the first comprehensive analysis of *KEAP1* mutations in thyroid tumors, revealing that these alterations are far more prevalent and broadly distributed than previously recognized. Although prior reports have only identified a small number of *KEAP1* mutations in PTCs and benign MNGs, we identified 81 such alterations across a diverse spectrum of thyroid cancers. Remarkably, we identified *KEAP1* mutations in all thyroid cancer histologies, ranging from well-differentiated forms (PTC, FTC) to PDTC and ATC, with no subtype specific mutational pattern. We also report the first instance of *KEAP1* mutations occurring in pediatric thyroid tumors. These KEAP1/NRF2 pathway alterations occurred at a frequency of 3.1% in our pediatric cohort, as compared to 0.6% in adult PTCs (TCGA), 2.0% in adult OTC (PRJNA445446), and 2.7% in adult PDTC/ATC (GSE76039).

In both pediatric and adult thyroid tumors, we identified *KEAP1* mutations with concurrent MAPK driver mutations, indicating potential cooperation between NRF2 and MAPK pathways in thyroid tumorigenesis. This group of dual *KEAP1-* and MAPK-mutant cases represents ~25% of all identified *KEAP1-*altered samples. We also identified a distinct subset of cases harboring both a *KEAP1* mutation and loss of heterozygosity (LOH) at chromosome 19p13.2, revealing a concerted genetic mechanism by which both alleles of *KEAP1* are inactivated through combined mutation and chromosomal loss. This biallelic loss occurred at a strikingly high frequency (~50% of cases with copy-number data), consistent with the classical two-hit model that underlies inactivation of many tumor suppressors. Interestingly, biallelic loss of *KEAP1* was mutually exclusive with MAPK alterations, suggesting two distinct modes by which *KEAP1* may promote tumorigenesis. In tumors harboring initiating MAPK driver alterations, activation of the NRF2 pathway via heterozygous *KEAP1* mutation may act as a cooperative event, enhancing tumor progression. Conversely, in tumors lacking MAPK mutations, biallelic inactivation of *KEAP1*, through combined single copy mutation and 19p13.2 LOH, may act as an initiating oncogenic event. Indeed, we identified seven cases of advanced disease (PDTC and ATC) and six PTC cases with no known driver mutations, but with full inactivation of *KEAP1*, supporting the theory that homozygous *KEAP1* loss may act as an oncogenic driver. These previously unclassified tumors highlight the importance of re-evaluating historically “driverless” cases with updated genomic tools. As sequencing technologies advance, our ability to uncover cryptic or novel oncogenic drivers – such as *KEAP1* inactivation – and characterize these “dark matter” tumors continues to expand. Although the majority of identified *KEAP1-*mutant cases lacked known driver mutations, limited CN data prevented us from determining whether these tumors harbored biallelic *KEAP1* loss. More comprehensive genomic profiling is essential to further examine the importance of biallelic *KEAP1* loss in thyroid tumors, particularly in cases that are genetically ambiguous.

For 78% of the cases in this report, we were able to confirm the germline or somatic status of the identified *KEAP1* alteration. Only five cases had confirmed germline alterations, three of which were benign multinodular goiters. The presence of these mutations is consistent with prior reports implicating *KEAP1* germline variants in familial multinodular goiter (44, 45, 69). Notably, the vast majority (92%) of alterations were somatic, further suggesting that *KEAP1* mutations may be functionally relevant in tumorigenesis. Although we recognize that some of the somatic *KEAP1* mutations listed may be passenger alterations, nearly half (48%) were predicted to be oncogenic or likely oncogenic by OncoKB. Evaluation of NRF2 pathway activation can help better profile the functionality of these mutations. Indeed, in 6 out of 7 cases where expression data was available, we observed upregulation of NRF2 target genes, indicating loss-of-function alterations. Notably, two of these mutations were not predicted by OncoKB to be oncogenic, but still showed significant NRF2 pathway activation, underscoring the value of a multifaceted approach to more accurately assess the pathogenicity of these mutations. The single *KEAP1* mutation lacking evidence of NRF2 pathway activation was a germline variant, co-occurred with a *CCDC186::RET* fusion driver mutation, and was predicted to be functionally neutral by SIFT, all supporting its likely status as a non-functional variant. While this MAPK-altered case lacked a functionally significant *KEAP1* mutation, the case carrying concurrent *KEAP1* and KRAS p.G12V mutations had robust NRF2 pathway activation, supporting that the mutation is functionally significant. These findings provide further support to the idea that in certain molecular contexts, NRF2 pathway activation may cooperate with MAPK mutations to enhance tumor progression. Notably, the co-occurrence of *KRAS* and *KEAP1* mutations has been linked to poorer clinical outcomes and reduced therapeutic efficacy in non-small cell lung cancer, further supporting a potential cooperative role of *KEAP1* alterations and MAPK activation (70). The strong correlation between NRF2 target gene expression and loss-of-function *KEAP1* mutations suggests the ability to infer *KEAP1* mutational status via an mRNA expression signature when accompanying DNA sequencing data is absent. Indeed, we identified four additional tumors with elevated NRF2 pathway activation in our transcriptomic analysis. While one case harbored a known *PAX8::NFE2L2* fusion, causing significant overexpression of NRF2 and its target genes, the remaining three lacked mutational profiling. Despite the absence of sequencing data for these tumors, their expression signatures showing NRF2 pathway activation strongly suggest underlying *KEAP1* inactivation, further emphasizing to the importance of paired genomic and transcriptomic information to determine both mutation presence and functional consequence.

We also report the first functional characterization of *KEAP1* mutations in thyroid cancer using *in vitro* models. To model the consequences of *KEAP1* loss both independently from and concurrently with MAPK driver alterations, we generated clonal *KEAP1* knockout cell lines from both TPC1, which harbors a *CCDC6::RET* fusion, and SDAR2, which lacks a known driver. In line with the patterns observed in patient tumors, we confirmed that *KEAP1* loss indeed induces NRF2 target gene expression in an NRF2-dependent manner in both MAPK-mutant and MAPK-wildtype cells, validating the strong association between *KEAP1* inactivation and NRF2 pathway activation. Consistent with the potential role of *KEAP1* mutations as novel oncogenic drivers, we found that loss of *KEAP1* promoted both cell proliferation and migration. These effects were observed in both cell lines, directly showing that NRF2 pathway activation via *KEAP1* loss can act independently or cooperate with MAPK alterations to drive tumor progression. These findings further implicate *KEAP1* mutations as functionally significant drivers in thyroid cancer, directly show their role in promoting tumorigenic cellular phenotypes across distinct genetic backgrounds, and position them as a potential therapeutic target in this disease.

In other cancers, overactivation of the NRF2 pathway has been shown to confer chemotherapeutic resistance; however, this resistance has not been explored in the context of MAPK inhibition (28, 33, 71). Here, we report the first instance of *KEAP1* mutations attenuating drug response in thyroid tumors. We show that loss of *KEAP1* reduces sensitivity of *RET* fusion-positive PTC cells to the receptor tyrosine kinase inhibitor selpercatinib, a key therapeutic agent for *RET*-driven thyroid cancers. These findings suggest that *KEAP1* inactivation may represent a previously unrecognized mechanism of resistance to *RET* inhibition in thyroid tumors. Our novel *in vitro* findings highlight the clinical relevance of *KEAP1* mutations in thyroid cancer and underscore the importance of mutational profiling to better inform therapeutic strategies.

Our comprehensive report of *KEAP1* mutations in thyroid tumors highlights its previously unappreciated prevalence and broad distribution across thyroid cancer subtypes. By combining multi-omic analyses with functional characterization of *KEAP1* loss in thyroid cancer, we demonstrate that these mutations may represent novel oncogenic drivers, capable of enhancing tumorigenesis across diverse genetic landscapes. Additionally, we identify a highly prevalent and concerted genetic mechanism in which both copies of *KEAP1* are lost. This two-hit mechanism, which is mutually exclusive with classical MAPK driver alterations, may represent the initiating oncogenic event in some tumors. This potential dual role – as either an initiating event or cooperating alteration – highlights the previously unrecognized importance of *KEAP1* mutations in thyroid cancer. Moreover, our finding that *KEAP1* mutations not only act as driver alterations but also attenuate cellular response to selpercatinib in *RET* fusion-positive cells, highlights their clinical significance. Our findings indicate that *KEAP1* mutations play an important role in driving tumorigenesis and modulating therapeutic response, suggesting that profiling of this pathway should be integrated into future studies and clinical testing.

## DATA AVAILABILITY

The data presented in this study are available from the corresponding author (ATF) upon reasonable request.

## ETHICS STATEMENT

This study was approved by Children’s Hospital of Philadelphia Institutional Review Board (IRB# 17-014224 and IRB# 20-018240). A waiver of consent was approved under certain conditions for existing biospecimens. Otherwise, written informed consent and assent to participate in this study were obtained from parents/guardians and/or participants.

## AUTHOR CONTRIBUTIONS

NEB, JCRF, and ATF designed the study. NEB, JCRF, ERR, CHP, KH, and AI performed the experimentation and acquired the data. NEB, JCRF, AJB, and ATF analyzed and interpreted the results. NEB, JCRF, and ATF drafted the manuscript, and NEB had primary responsibility for final content. All authors contributed to the article and approved the submitted version.

## CONFLICT OF INTERSET

The authors declare that the research was conducted in the absence of any commercial or financial relationships that could be construed as a potential conflict of interest.

## ACKNOWLEDGEMENTS

We would like to acknowledge Zachary Spangler, Michele Scheerer, Kay Labella, and the entire Franco Laboratory for their support throughout the duration of this study and preparation of this manuscript.

