## Supplemental Figures for "*KEAP1* mutations activate the NRF2 pathway to drive cell growth and migration, and attenuate drug response in thyroid cancer"

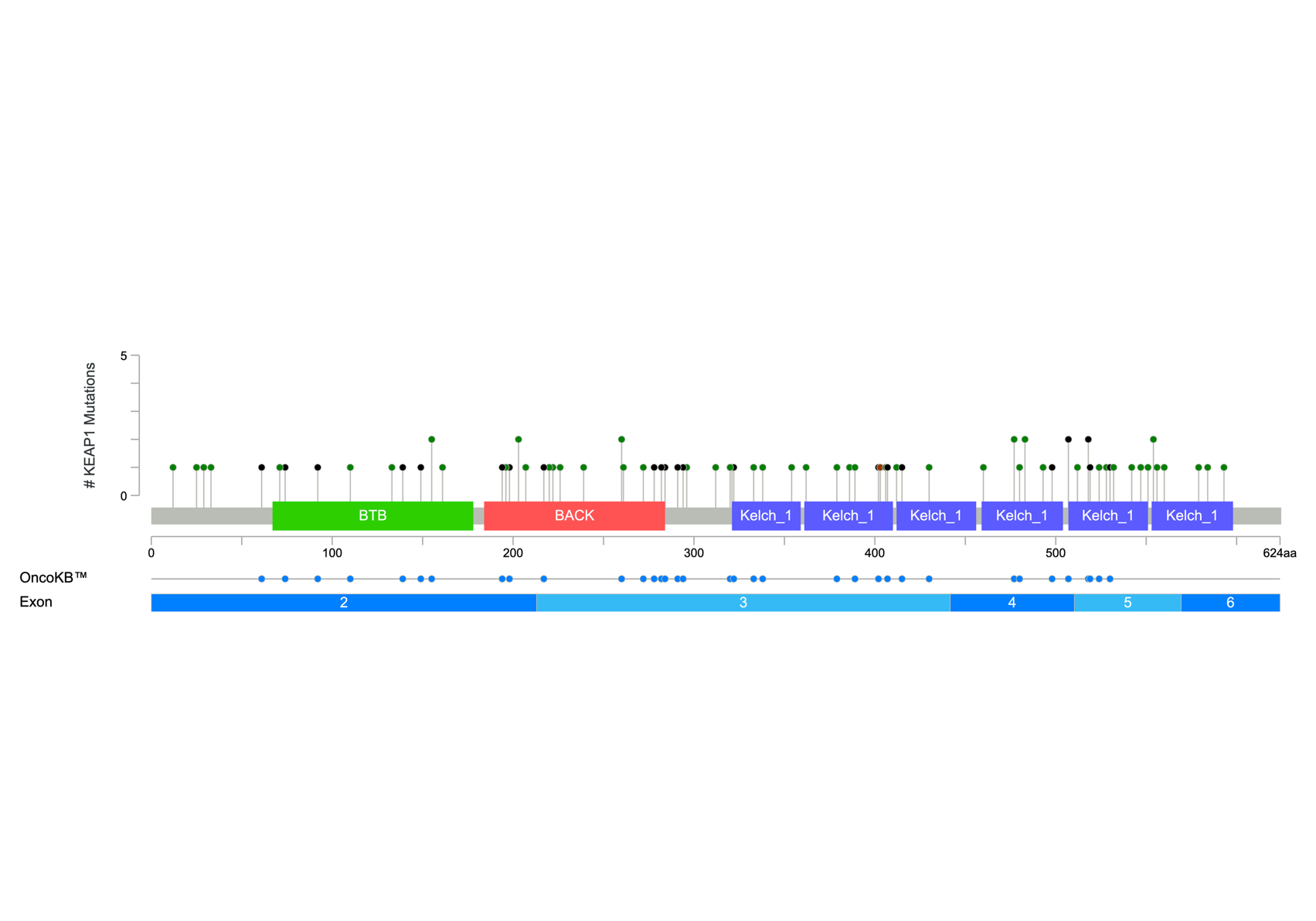
**Supplementary Data**

**Figure S1.** Mutation mapper showing distribution of the 81 *KEAP1* mutations across gene exons and annotated protein domains. Green dots represent missense variants, black dots represent truncating variants, and brown dots represent in-frame deletions. Variants are annotated based on their OncoKB predictions, with blue dots representing predicted pathogenic mutations.

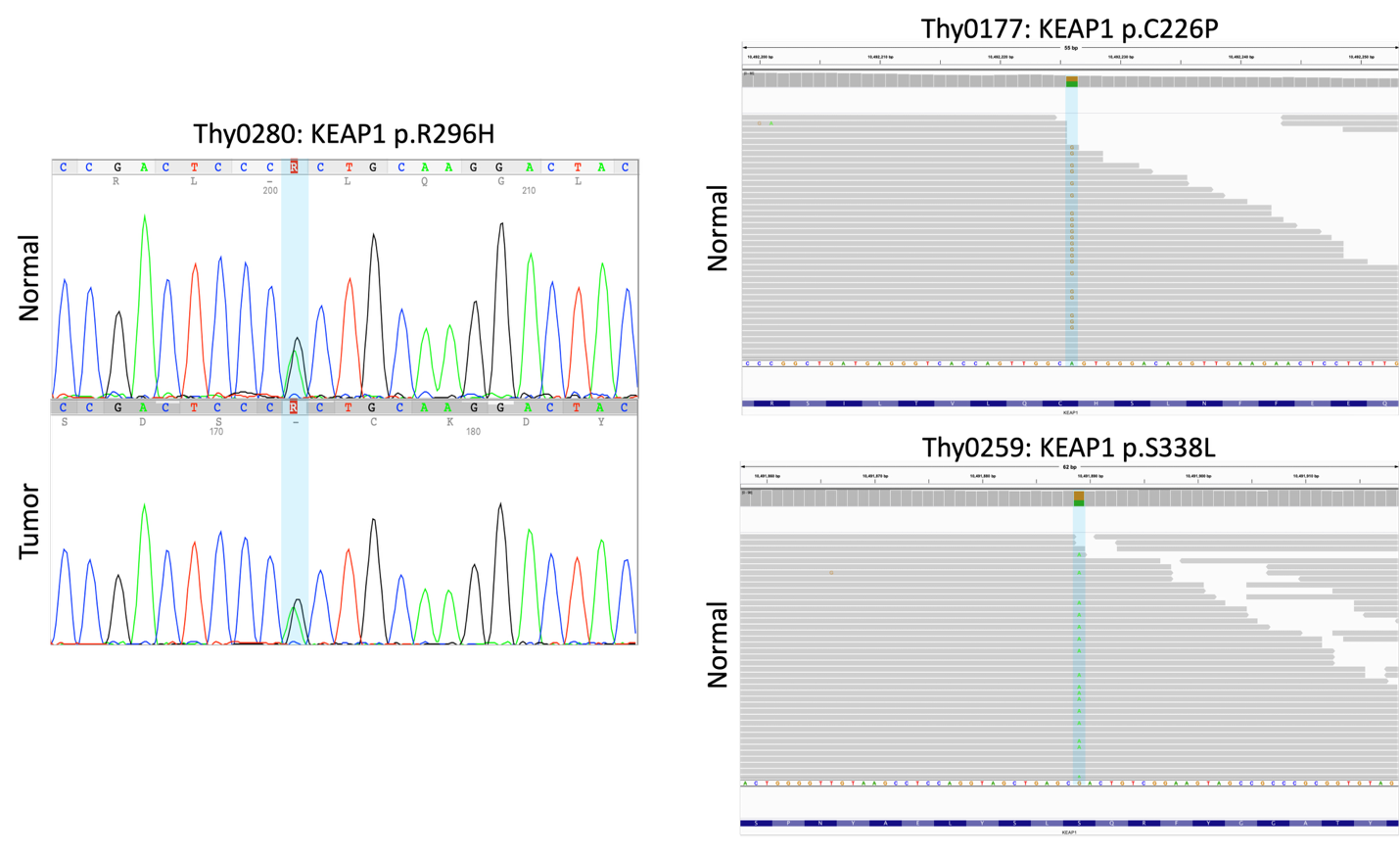

**Figure S2. Evaluation of *KEAP1* germline status in CHOP cohort samples.** Sanger sequencing traces of Thy0280 tumor and adjacent normal samples show germline heterozygous status of KEAP1 p.R296H alteration. IGV visualization of aligned whole exome sequencing reads for Thy0177 and Thy0259 adjacent normal tissue confirms germline heterozygous status of KEAP1 p.C226P and KEAP1 p.S338L mutations, respectively. Alterations are highlighted in blue.

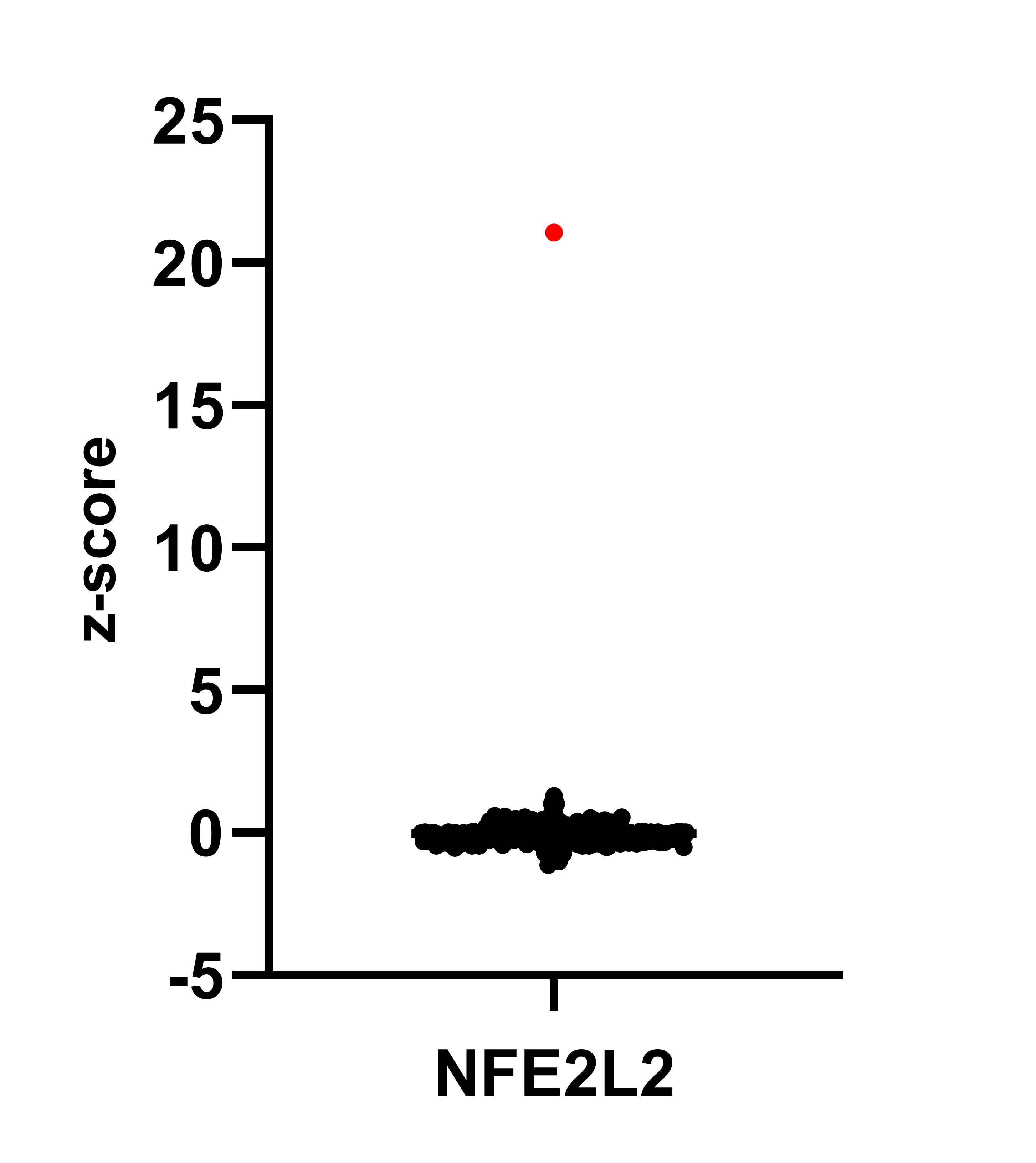

**Figure S3.** RNA expression of *NFE2L2* for samples in the TCGA cohort demonstrate that the *PAX8::NFE2L2* fusion-positive case (red dot) exhibits significantly higher expression of *NFE2L2* (gene encoding NRF2 protein)*.* Z-scores were calculated relative to all samples in the cohort.

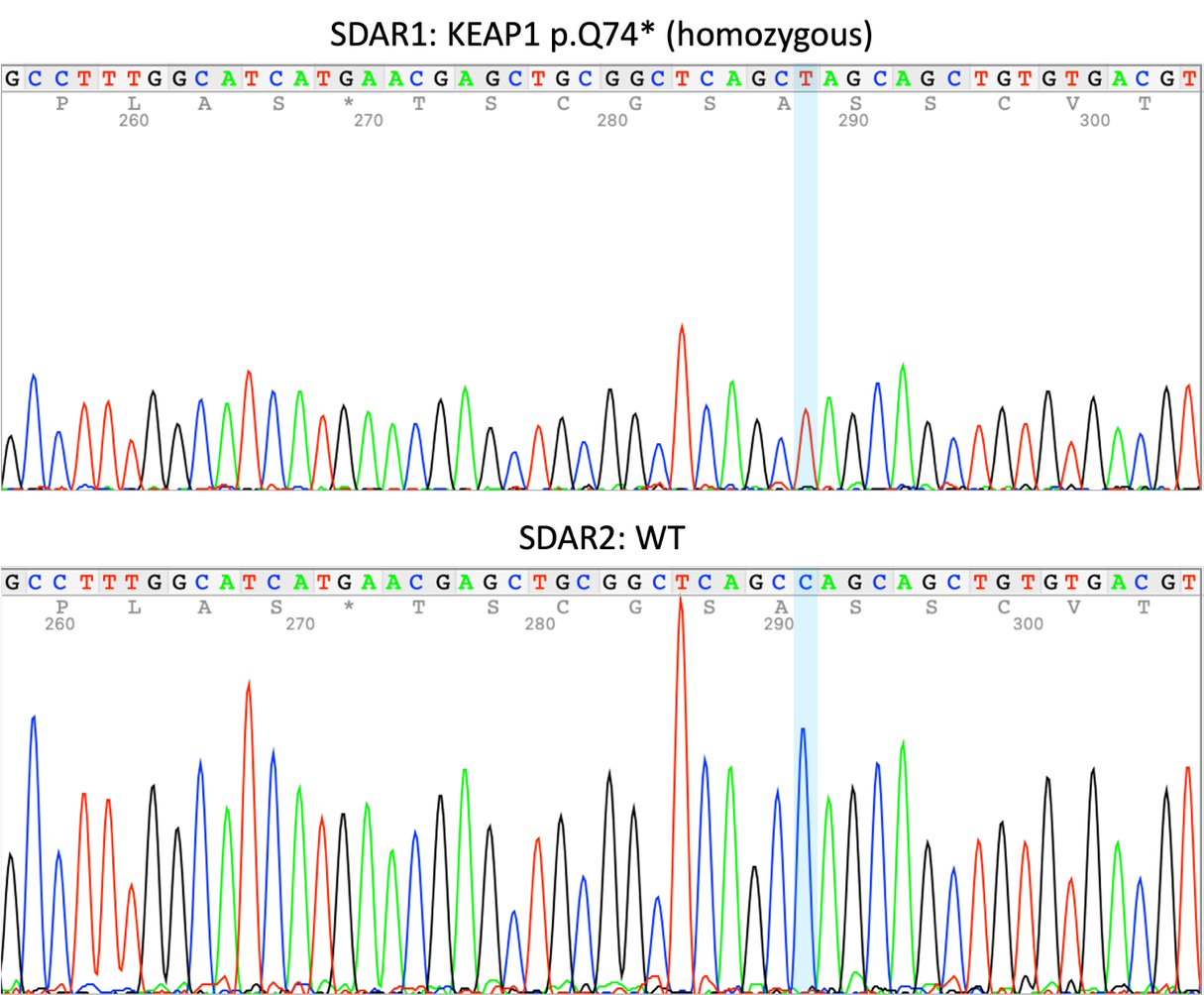

**Figure S4. Validation of SDAR1 *KEAP1* nonsense mutation.** Sanger sequencing of exon 2 of *KEAP1* shows a homozygous *KEAP1* c.220C>T (p.Q74*) point mutation in SDAR1, while SDAR2 retains both wildtype alleles. The alteration is highlighted in blue.

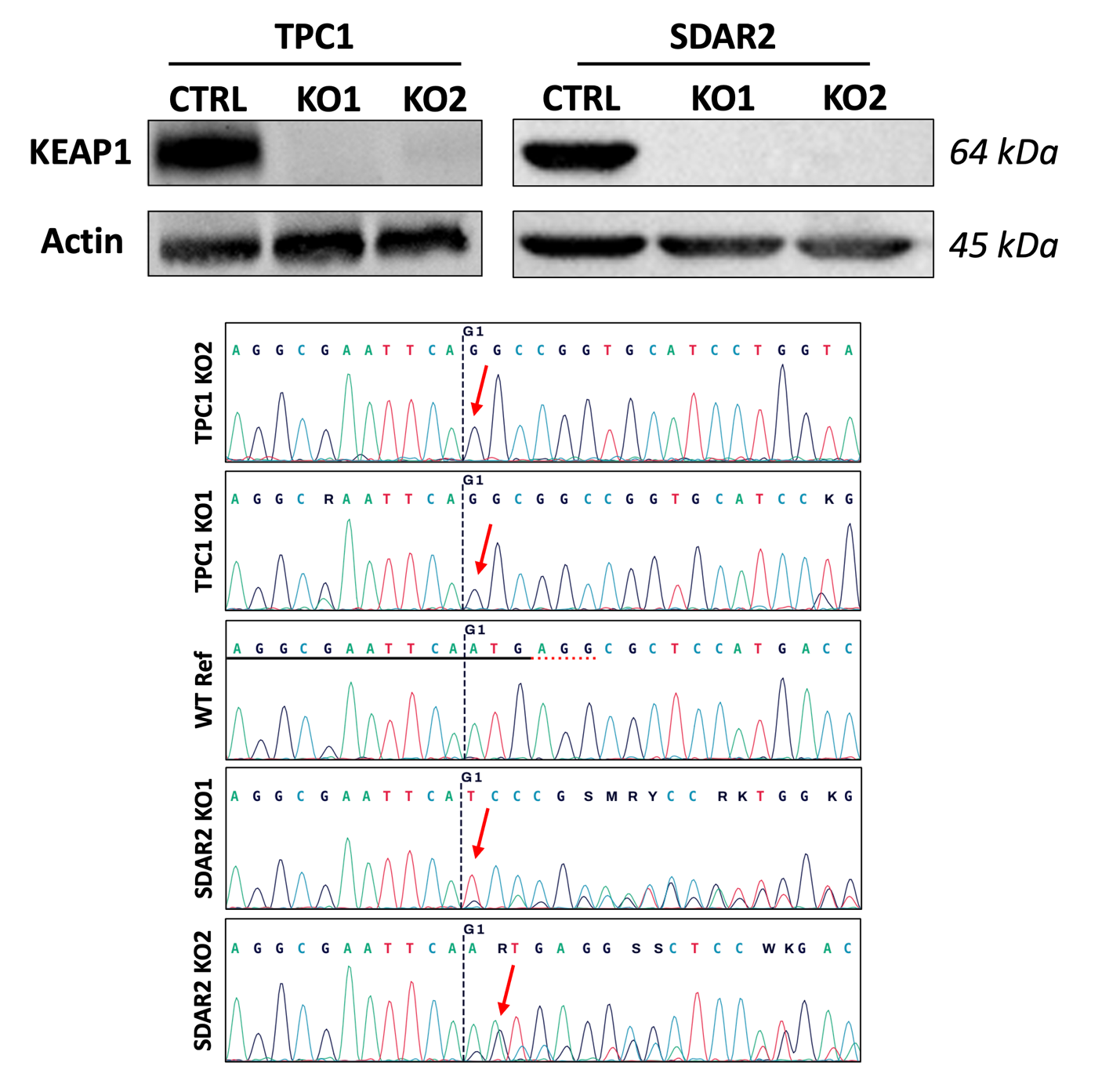

**Figure S5. *KEAP1* knockout cell line validation.** Western blotting confirms the absence of KEAP1 protein in both the TPC1 and SDAR2 knockout clones. β-actin was used as a loading control. Sanger sequencing was performed on PCR-amplified fragments containing sgRNA binding sites in both TPC1 and SDAR2 knockout clones, as well as a wildtype reference. Sequencing shows successful DNA-level alterations in both TPC1 and SDAR2 knockout clones. The cut site for sgRNA1 is shown as a black dashed line and red arrows highlight alterations in the knockout clones occurring around this site. The sgRNA binding region and PAM site is shown in the WT Ref trace as a solid black line and dashed red line, respectively.
