## Supplemental Materials and Methods for "*KEAP1* mutations activate the NRF2 pathway to drive cell growth and migration, and attenuate drug response in thyroid cancer"

**Supplementary Materials and Methods**

**Sanger sequencing primers**

All sequencing primers were designed using NCBI Primer-Blast to generate products of approximately 300-400 bp. Primers for *KEAP1* knockout validation were designed to amplify the region containing all three guide RNA cut sites. Primers for mutation confirmation were designed to amplify the known point mutation. The following primers were used:

| **Primer** | **Sequence** |
| --- | --- |
| KEAP1 KO VALID FWD | GTAGATGTACTCCCGGGCAC |
| KEAP1 KO VALID REV | CAACCGCACCTTCAGCTACA |
| THY0280 R296H FWD | GTCACTGGGGTTGTAAGCCT |
| THY0280 R296H REV | TCTTCAACCTGTCCCACTGC |
| SDAR Q74* FWD | CCATCTCCATGGGCGAGAAG |
| SDAR Q74* REV | GTAGATGTACTCCCGGGCAC |

**RT-qPCR primers**

All primers were designed using NCBI Primer-Blast and were positioned to span introns as to avoid amplification of DNA contaminants. The following primers were used:

| **Primer** | **Sequence** |
| --- | --- |
| Hs_NQO1_qPCR_Fwd | AGGACATCACAGGTAAACTGAAG |
| Hs_NQO1_qPCR_Rev | TATCACAAGGTCTGCGGCTT |
| Hs_AKR1C3_qPCR_Fwd | AAAAAGCACAAGCGAACCCC |
| Hs_AKR1C3_qPCR_Reve | TCCTCTGCAGTCAACTGGAAC |
| Hs_GCLC_qPCR_Fwd | AAAAAGTCCGGTTGGTCCTGT |
| Hs_GCLC_qPCR_Rev | GTCCTGGTGTCCCTTCAATCA |
| Hs_TXNRD1_qPCR_Fwd | CCAGGACATGGCCAACAAAAT |
| Hs_TXNRD1_qPCR_Rev | TACTACTCTGAGTCGGCCTGG |
| Hs_ACTB_qPCR_Fwd | TACAATGAGCTGCGTGTGGC |
| Hs_ACTB_qPCR_Rev | AGCACAGCCTGGATAGCAAC |
